# The internal metabolic state controls behavior in *Hydra* through an interplay of enteric and central nervous system-like neuron populations

**DOI:** 10.1101/2023.09.15.557876

**Authors:** Christoph Giez, Christopher Noack, Ehsan Sakib, Thomas Bosch

## Abstract

Hunger and satiety can have an influence on decision making, sensory processing, and motor behavior by altering the internal state of the brain. This process necessitates the integration of peripheral sensory stimuli into the central nervous system. Interestingly, even organisms without a brain, such as the Cnidaria, exhibit feeding dependent behavioral changes. The underlying mechanisms, however, remain unclear. In this study, we demonstrate that neuronal activity in two distinct neuronal populations, the ectodermal N3 neurons and the endodermal N4 neurons in *Hydra*, an ancestral metazoan animal with a diffuse nerve net spread throughout the body with no signs of centralization, are responsible for feeding dependent behavioral changes. Specifically, endodermal N4 neurons are essential for food intake and digestive functions, similar to the enteric nervous system, while the N3 population influences and inhibits other motor related behaviors, comparable to the central nervous system. This fascinating observation provides a new insight into the evolution and the complexity of a simple non-centralized nervous system.

## Introduction

Most if not all animals exhibit behavioral responses in relation to food consumption or starvation. Satiety and hunger are two metabolic states which have a global effect and influence on decision making, sensory processing and motor behavior^1–3^. In the majority of animals including human, the integration of such information occurs in the brain which then effects the outcome of behaviors. The signal of satiety emerges in the periphery, in the gastrointestinal tract and enteric nervous system^4^. Interestingly, internal states such as hunger or satiety can be already observed in organisms without a centralization of the nervous system and an enteric nervous system such as in the phylum Cnidaria^5–8^. Cnidaria (i.e., sea anemones, jellyfish, corals and hydra) and Bilateria (i.e., vertebrates, sea stars, fruit flies), are sister groups that diverged around 600 million years ago (**Fig. 1A**) ^9^. In the jellyfish *Clytia*, starvation had an influence on the food passing behavior^5^. In *Cassiopea* and *Hydra* a sleep-like state has been observed where they exhibit phases of quiescence^6,7^. Further, *Hydra* seems to have a feeding state dependent phototaxis behavior where it is more attracted to light when starved^8^. However, the neuronal basis of those internal states has not been elucidated so far.

**Figure 1.**
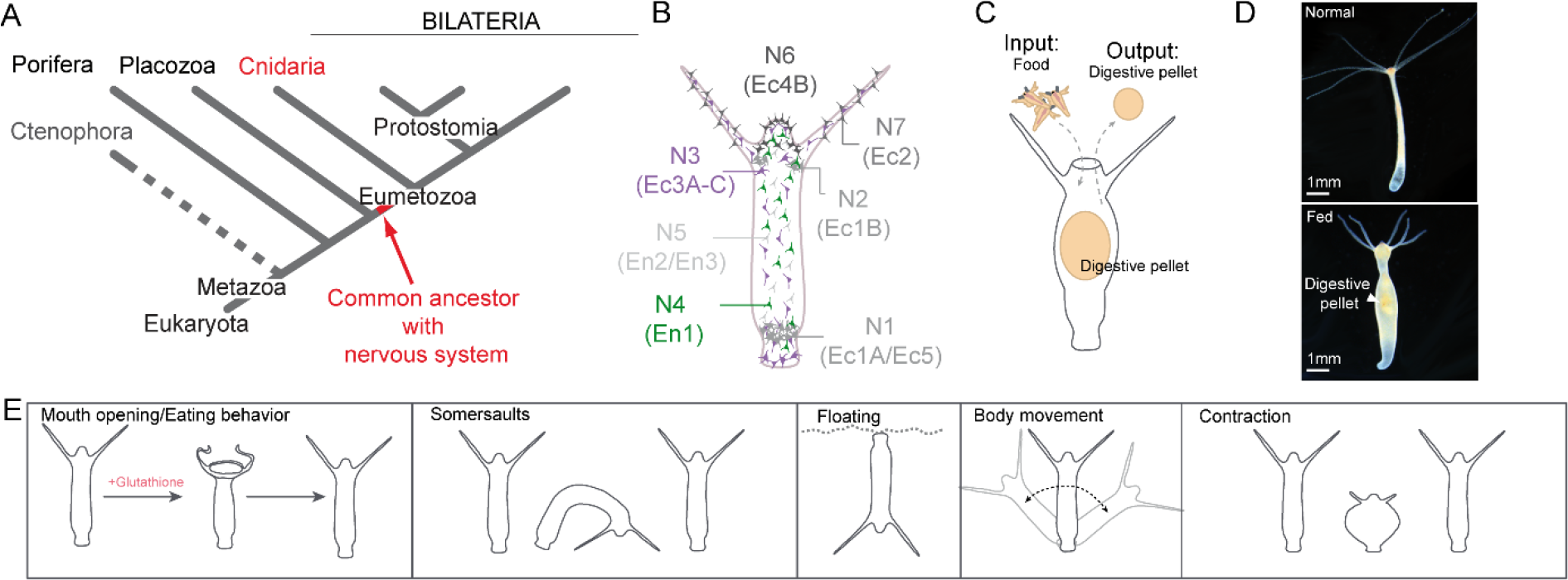
*Hydra* as a model system to study the evolution of the nervous system. **A.**Phylogenetic positioning of *Hydra* at the origin of the nervous system with the same building tools than those of Bilaterian. **B.**The different neuronal populations identified by single cell sequencing^33–35^. The different nomenclature highlighted here. **C.**Input of food and output of remaining digestive pellet is the same in *Hydra*. **D.***Hydra* not fed and *Hydra* 4 hours post feeding. **E.**Different behavioral pattern in *Hydra*: Mouth opening due to glutathione, somersaults, floating at the surface, body movement or swaying and contractions.

The tubular bodies of Hydra are radially symmetric and have only two layers, an ectodermal and an endodermal epithelium. A gastrovascular cavity has a single exterior opening that serves as both mouth and anus. Tentacles surround the mouth opening. Nerve cells are organized into two simple nerve nets, one ectodermal and the other endodermal, consisting of multiple neuronal populations that help coordinate muscular and sensory functions (**Fig. 1B - D**). Previous work has shown that the eating behavior of *Hydra* can be induced by reduced glutathione ^10,11^. Interestingly, after one GSH-induced full eating behavior a recovery period of over 24h has been described^10^. However, despite this, the polyps were still able to ingest living prey ^12^. The sensitivity to glutathione seems to depend on the feeding status since well-fed animals did not exhibit a feeding response^10,13^. Those papers already speculated about an internal state suggested to control the eating behavior ^14^.

Here we explored the possibility of satiety dependent behavioral changes in *Hydra* and the neuronal mechanism underlying the satiety-driven modifications that integrate information related to feeding status. Additionally, we examined whether there is a sub-functionalization of the nervous system regarding feeding-related behaviors and another for motor and information processing. Our study shows that *Hydra*’s behavior changes significantly depending on the degree of satiety. Immediately after the animal has taken food, *Hydra* stops contracting, somersaulting and shows no response to glutathione (food stimulus) but an increase in body movement (**Fig. 1E**). This drastic change in behavior arises from the ectodermal neuronal network N3 which shows a feeding dependent activity change. The genetic ablation of N3 neurons leads in well fed animals to a decrease in body movement, increased phototaxis and to a longer mouth opening induced by glutathione. The endodermal neuronal network N4 responds differently depending on the presence and absence of a digestive pellet in the gastric cavity. Ablation of enteric-like N4 neurons leads to a bursting of the entire well-fed polyp beneath the hypostome since the animals lost the ability to spit out the digestive pellet. In summary, we identified the ectodermal neuronal network N3 as being responsible for encoding the metabolic internal state and an endodermal neuronal network N4 which is necessary for feeding and digestion associated behaviors. These findings may reflect an early separation of a diffuse neural network into a population of centralized and population-cooperating neurons and an enteric nervous system-like population.

## Results

Ever since the eating behavior of the classical model organism *Hydra* has been described, multiple papers have reported that there is a refraction time or habituation to the stimulus of glutathione. Here, we did a first thorough investigation of this phenomenon and discovered that satiety of *Hydra* has a global effect on multiple behaviors but does not depend on glutathione exposure.

### Satiety changes behavioral patterns in *Hydra*

To show that an exposure to glutathione or the extensive feeding of *Hydra* have an impact on the re-exposure to glutathione, we tested animals for their eating behavior which were either extensively fed with *Artemia* or exposed to glutathione (chemical food signal) for 2 hours. Animals which were extensively fed with *Artemia* showed only little response to glutathione until 10 hours post feeding while having normal behavior after 24 hours again (**Fig. 2A**). In a similar manner, their response time (the time until the mouth opens) was significantly slower within the first 8 hours post feeding while at 10 hours they responded normally fast again (**Fig. 2B**). In contrast, animals exposed to glutathione for 2 hours prior to the experiment showed a normal behavior after 4h post exposure (**Fig. 2A-B**). However, their response time was slightly slower over a longer period of time (**Fig. 2B**). In summary, only feeding *Artemia* resulted in a sensation of satiety with subsequent and long-lasting changes in mouth opening/eating behavior. Exposure to glutathione had only small effects.

**Figure 2.**
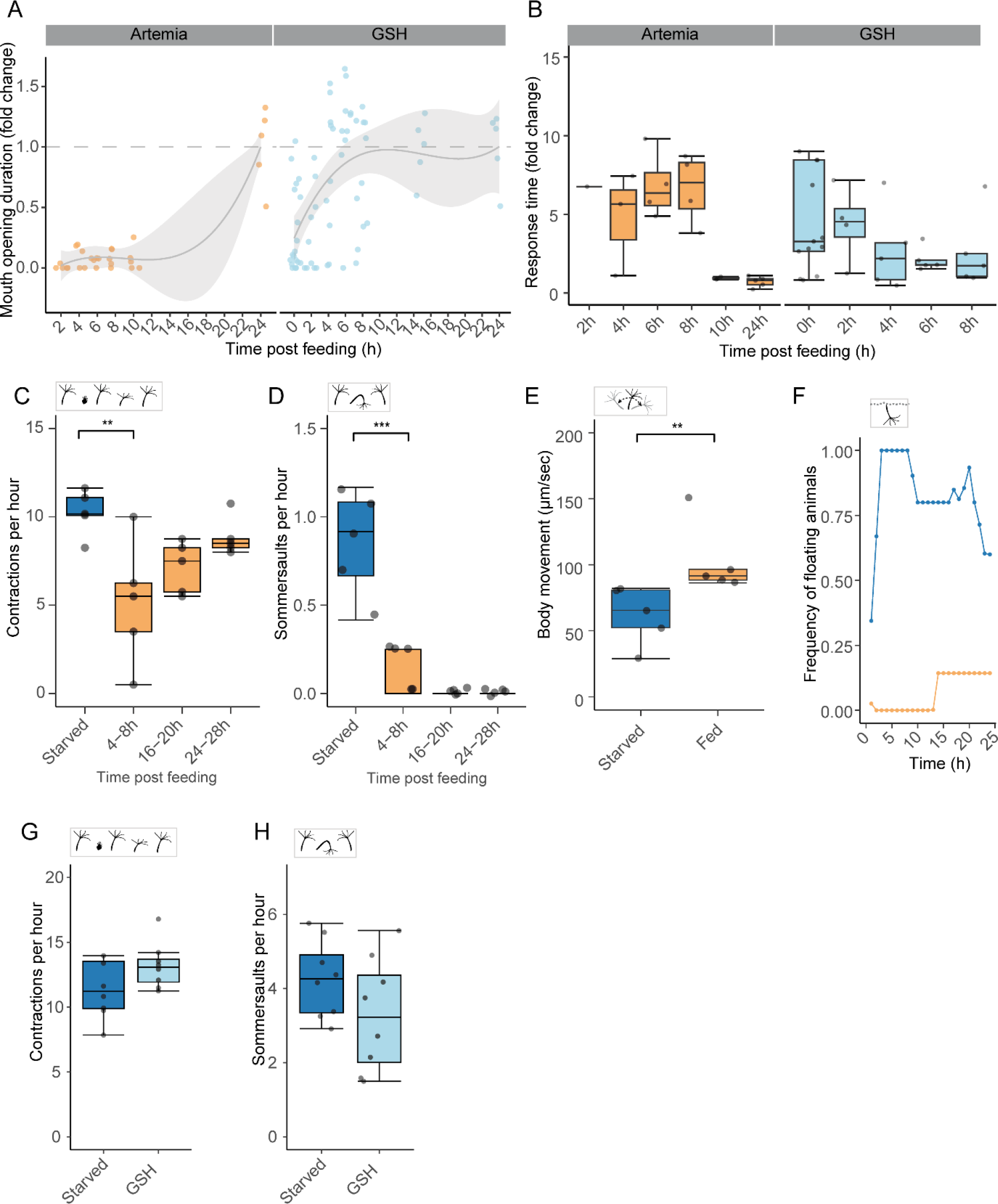
Satiety has a global impact on different behaviors and depends on the ingestion of digestible food. **A.**After feeding animals extensively, the feeding response cannot be induced by reduced glutathione (GSH) till 10 hours post feeding (orange). The feeding cannot be mimicked by incubating animals for 2h in GSH prior experiment (light blue). n = 5-10 **B.**After feeding the response time to GSH is delayed for 8 hours post feeding whereas the effect of GSH exposure vanishes much faster (n= 5-10). **C.**Comparing starved animals (10 days without feeding) to fed animals, a difference in contractions per hour was observed for 4-8 hours post feeding while afterwards the frequency returns to the starved conditions (n= 5, ANOVA, t-test, p<0.01). **D.**Somersaults were almost absent in fed animals whereas starved animals made 1 somersault per hour (n =5, Kruskal-Wallis test, Dunn test, p<0.001). **E.**The movement of the body center by any kind of direction change was slightly increased in fed compared to starved animals (n = 5, Kruskal-Wallis test, Dunn test, p<0.01). **F.**Starved animals had a higher proportion of animals floating (detached and at the water surface) than fed animals (n = 5). The effect persisted over 24h. **G-H**. The effect on contractions and somersaults could not be affected by pre-exposure to GSH (2h incubation). (n = 5)

Observing the effect of satiety on the eating behavior, we were curious if other movement patterns of *Hydra* such as spontaneous body contractions, somersaults, or movements in general were also affected. Therefore, we next explored a range of behaviors of animals after being fed and scored their behavioral patterns (**Fig. 1E, Fig. 2C-F**). We observed that contractions were significantly reduced (p<0.01, n = 5) until 8 hours post feeding but increased thereafter and reached the contraction frequency of starved animals again around 24 hours post feeding (**Fig. 2C**). In case of somersaults, fed animals displayed a notable absence of somersaults which was significantly less than starved animals (p< 0.001, **Fig. 2D**). Interestingly, measurements of movement from the center of the body (see **Fig. 1E**) point to large overall movements of the polyps (**Fig. 2E**). Given the absence of observed somersault events, next we wanted to find out if starved animals are more frequently detached and float on the surface. By comparing the proportion of floating animals between starved and fed animals, we examined that starved animals were mainly floating whereas fed animals did not show any detachment until 13 hours post feeding (**Fig. 2F**). The behavioral changes were specific to animals which were fed with *Artemia* since the same behavioral changes could not be observed with glutathione exposure (**Fig. 2G-H**). In summary, satiety has a global and long-lasting effect on the behavior of the polyp *Hydra*. The stimulus responsible for this clearly comes from digestion and engulfment of real food and cannot be replaced by glutathione.

### Neuronal populations 3 and 4 respond to satiety very differently

Intrigued by the behavioral changes, next we investigated if the effects are induced by changes in neuronal activity. To approach the nature of the neuronal control of satiety in *Hydra*, we conducted a systematic screening of different neuronal populations by using previously established calcium imaging lines. In a first step, we took the advantage of a pan-neuronal *Hydra* line (alpha-tubulin promotor^15^) and investigated neuronal activity in the body column (**Fig. 3A**). We observed a significant increase in neuronal activity in the body column when comparing starved and 4 hours post feeding animals (p<0.01). However, it was not possible to distinguish between distinct neuronal subgroups and to identify the specific population accountable for the observed phenomenon. Therefore, we explored all different neuronal networks which were described so far (**Fig. 1 B, Fig. 3C**) ^16–18^. First, we examined the N1 neuronal network - previously named contraction burst (CB) - which was correlated with body contractions ^17^. The N1 network showed a slight decrease in its frequency in fed animals (**Fig. 3B**). Second, we analyzed the spiking frequency of the endodermal neuronal population N4 (**Fig. 3C and F**). The N4 neuronal displayed no notable distinction between animals which were fed and had expelled their digestive pellet compared to starved animals (**Fig. 3D**). Most interestingly however, animals that had not expelled their pellet and still retained it in the gastric cavity behaved quite differently by showing a significant increase in their spiking frequency (p< 0.05, **Fig. 3D**). Finally, we analyzed the spiking frequency of the ectodermal neuronal population N3 (**Fig. 3C and I**). N3 neurons showed a significant increase in spiking, independent of the presence or absence of a digestive pellet in the gastric cavity when compared to starved animals (p< 0.01, **Fig. 3G**). Since a change in the mouth opening duration has been observed in fed animals (**Fig. 2A-B**), we aimed to investigate whether there was a correlation between the significant difference in activity observed in the two populations, N3 and N4 (**Fig. 3D and G**), and the altered mouth opening duration. To accomplish this, we investigated the neuronal activity during mouth opening with a previously established calcium imaging setup^16^. We observed that the endodermal N4 and the ectodermal N3 neuronal population responded significantly different compared to the starved control (**Fig. 3E and H**). The N4 population displayed a reduced frequency whereas N3 showed an increased frequency during the mouth opening induced by glutathione.

**Figure 3.**
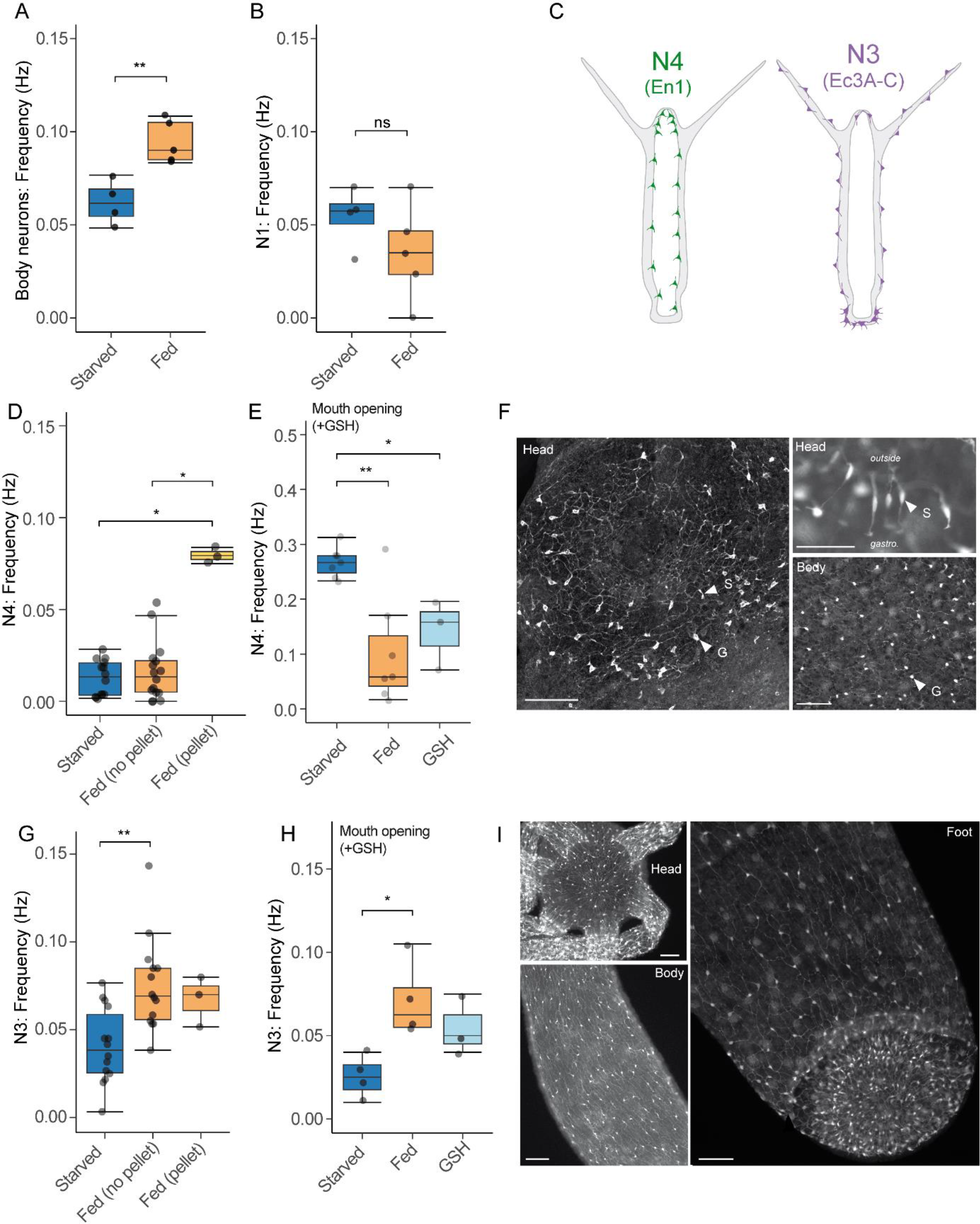
Two globally distributed neuronal networks respond differently 4 hours post feeding and do not respond normally to glutathione. **A.**Neuronal activity increased in the body column of a pan-neuronal line after 4 hours post feeding compared to starved animals. All neuronal activity was analyzed, no differentiation between populations (n = 5, T-test). **B**. Neuronal activity of the CB/N1 network in the foot of animals either starved or 4 hours post feeding showed no difference (n = 4-5, T-Test). **C**. Schematic of the two different neuronal populations in *Hydra*. N3 in the ectoderm and distributed along the whole-body axis. N4 in the endoderm and only distributed in the body column and head. **D**. Endodermal N4 neuronal population showed no difference 4 hours post feeding compared to starved animals when there is no pellet inside the animals. Animals with a digestion pellet showed a significant increase in spiking frequency. (n = 3-16, ANOVA, Turkey) **E**. Endodermal N4 population did not show an increase in spiking frequency as starved animals when exposed to reduced glutathione 4 hours post feeding or 1 hour post GSH exposure (n= 3-6, ANOVA, Turkey). **F**. Immunohistochemistry of endodermal neuronal population N4 in different locations of *Hydra*. In the head N4 has a spiderweb similar arrangement with sensory cells (S) and ganglion cells. *Scale bar 100µm* **G**. Ectodermal N3 population increased in spiking frequency 4 hours post feeding compared to starved animals. The presence or absence of a pellet did not affect the spiking frequency (n = 3-17, ANOVA, Turkey). **H**. Ectodermal N3 population did not decrease in spiking frequency compared to starved animals when exposed to glutathione 4 hours post feeding or 1 hour post feeding. (n= 3-6, ANOVA, Turkey) **I**. Immunohistochemistry of ectodermal population N3 at different locations in *Hydra*. Highest neuronal density is found in the foot. *Scale bar 100µm* p-values: * p<0.05, ** p< 0.01, *** p< 0.001

### Ectodermal N3 neurons control motor behavior

With the ectodermal N3 neuronal population displaying an increase in activity after feeding, we next wanted to determine the implications of this altered activity pattern. The N3 neuronal network is distributed all over the polyp with a higher density in the foot region and displays synchronous firing (**Fig. 3C and I**). Fed animals had an increase in spiking which showed a striking regular spiking pattern after 4 hours post feeding compared to starved animals (**Fig. 4A-B**). The spiking frequency decreased over time post feeding until it reached a frequency similar to starved animals, approximately around 15 hours post feeding (**Fig. 4C**). The activity of N3 did change in a behavior specific manner. Elongation behavior had a significant increase in spikes compared to starved animals (**Fig. 4D**). Interestingly, non-visible behavior associated spikes also increased significantly (**Fig. 4D**). Since N3 previously has been associated with locomotion behavior in *Hydra* ^17^, we hypothesized that N3 plays a major role in motor behaviors and their regulation in fed animals. The causal role of N3 neurons became clear when we discovered in an ablation experiment a highly significant decrease in body movement in fed transgenic polyps compared to wildtype fed animals (**Fig. 4E**). N3 has also been proposed to have an inhibitory impact on the eating behavior and the mouth opening ^16^. Therefore, we next tested, if the absence of N3 neurons affects the response time or the mouth opening duration in animals 4 hours post feeding. Interestingly, N3 ablated animals responded significantly faster to glutathione than the fed control (p< 0.01, fed wildtype with MTZ) and showed a significant increase in mouth opening duration (p< 0.05, **Fig. 4F-G**). In addition, N3 ablated animals exhibited multiple mouth opening events which could not be observed in the control (**Fig. 4G**). To show that a change in frequency underlies the effect, we reduced the spiking frequency of N3 by cutting off the foot of animals and then tested their response to glutathione. As a result of the absence of artificial methods to decrease the spiking frequency, we observed that the excision of the animal’s foot led to a reduction in the spiking frequency within the N3 population (**Fig. 4H**). In accordance with our predictions, cutting off the foot led to a significant increase of mouth opening duration in fed animals compared to starved ones (p< 0.01, **Fig. 4I**). In addition, we observed that the feeding dependent phototaxis behavior was changed in well fed N3 ablated animals. While normally an increased phototaxis behavior was observed in starved animals ^8^, N3 ablated animals showed an increased movement behavior towards the light source (**Fig. 4J**).

**Figure 4.**
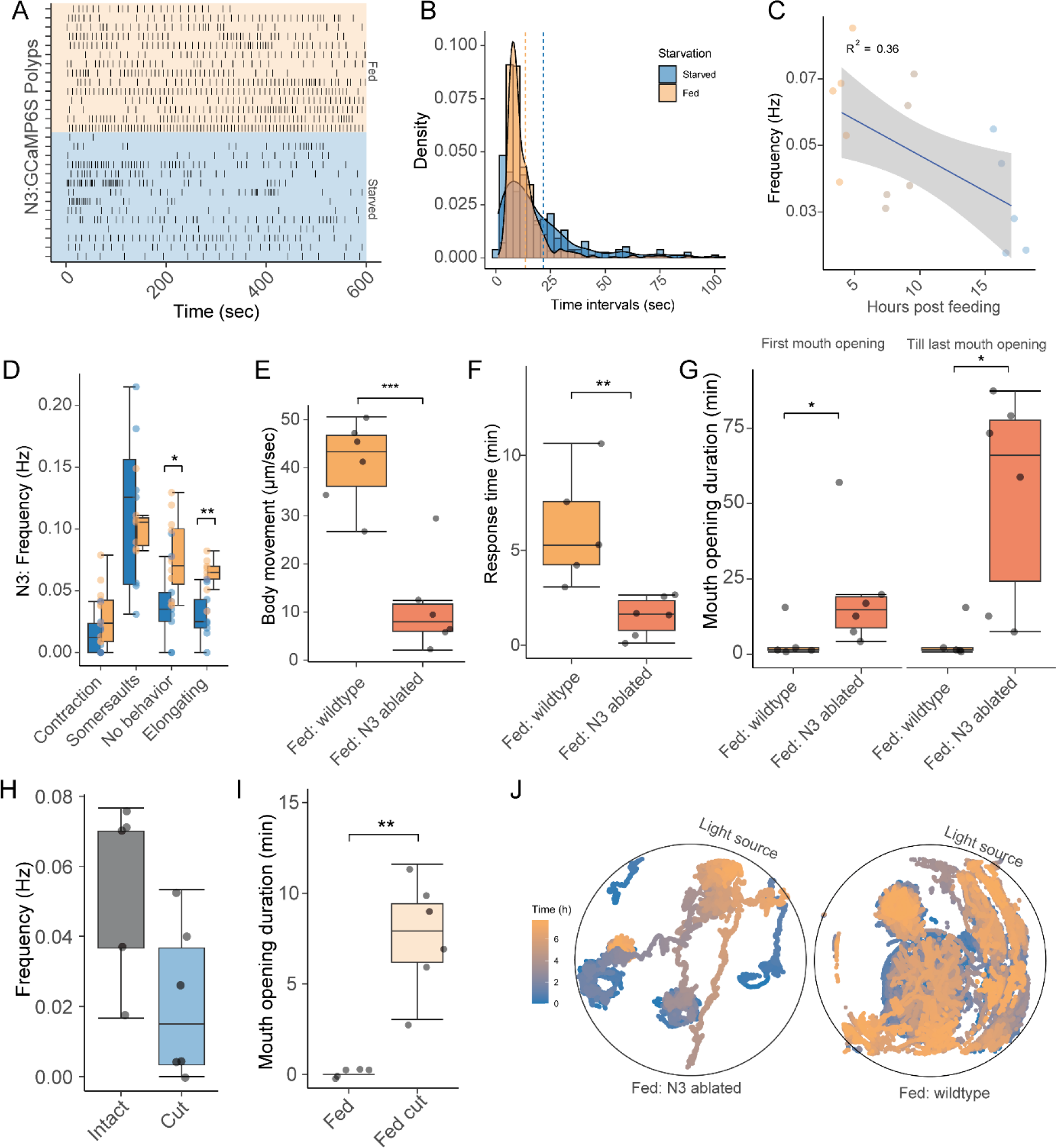
N3 neurons spikes regularly and prevents mouth opening as well as general body movement post feeding in a spiking frequency dependent manner. **A**. Spiking trains of either fed (4 hours post feeding) and starved animals. **B**. Inter spike intervals distribution across the two different treatments: starved and fed. More shorter and regular spikes observed in fed animals than starved. **C**. Ectodermal N3 population decreased in spiking frequency post feeding and showed a negative correlation (n= 5). Each data point responds to an independent animal post the respective time of feeding. **D**. N3 neurons exhibited different spiking frequency during different behaviors associated with N3. In fed animals (4h post feeding) the spiking frequency during elongation and non-behavior associated spiking is significantly increased. (n = 5-10, T-test) **E**. In well fed animals where N3 neurons were ablated, almost no body movement was observed, whereas wildtype animals exhibited higher mobility. (n=6-7, T-test) **F**. Ablation of N3 restored response time to glutathione of fed animals. Animals responded within 1.63±1.04 min to GSH whereas fed animals responded 10±35.8 min. (n = 5-6, Wilcox-Test) **G**. Ablation of N3 led animals respond to GSH for a longer period than fed animals. N3 ablated animals showed multiple mouth openings (≥2). (n = 5-6, Wilcox-Test) **H**. In animals without a foot the spiking frequency of N3 is decreased compared to intact polyps (n = 6). **I**. When exposed to glutathione animals without a foot respond again with a mouth opening (n= 6). **J**. Phototaxis behavior is elevated in N3 ablated animals, a behavior which is normally suppressed in fed animals. Tracks of six different is colored based on the time (blue: start, orange: end). (n = 6)

### Endodermal N4 neurons control food intake and the spitting out of the digestive pellet

Next we studied the N4 neurons and noticed that the spiking frequency between starved and fed animals without a digestive pellet is rather similar. Animals that still had a digestive pellet in their gastric cavity, however, showed increased spikes and shorter inter spike intervals (**Fig. 5A-B**). The endodermal N4 population is distributed along the whole-body axis except for the tentacles with an increase in density in the hypostome area (**Fig 3 C**)^16^. Within the body, the population forms a network composed of ganglion neurons whereas in the head it forms a spiderweb-like structure with ganglion and sensory neurons which extend into the gastrovascular cavity (**Fig. 3F**). First, we have tested whether or not animals with an ablated N4 population could continue to ingest their prey. As *Hydra* would not have been able to catch prey due to MTZ-treatment, we decided to use artificial food (here glass beads) for testing its ability to swallow the prey. Consistent with previously published data^16^, all N4 ablated animals could open their mouth. Surprisingly, however, they cannot swallow the glass beads and transport them into the gastric cavity (**Fig. 5C**). In contrast, the control group successfully engulfed all presented glass beads (**Fig. 5C**, control: wildtype + MTZ). N4 neurons also appear to be responsible for facilitating the digestive pellet to leave the gastric cavity. A significant time delay in spitting out glass beads was observed in N4 ablated animals, further supporting the role of N4 in the spit-out process (p< 0.05, **Fig. 5D**). We then investigated whether the expulsion time changes when using *Artemia* instead of glass beads. To our great surprise, although the expulsion time did not change **(Fig. 5E**), in N4 ablated polyps the pellet could not leave the gastric cavity via the normal exit, i.e. the mouth. Amazingly, fed animals without N4 neurons first became inflated like a balloon beneath the hypostome region, and then burst, releasing the digestive pellet through the ruptured tissue (**Fig. 5F and G**). N4 neurons, therefore, appear to be responsible for both controlling food intake and excreting the digestive pellet.

**Figure 5.**
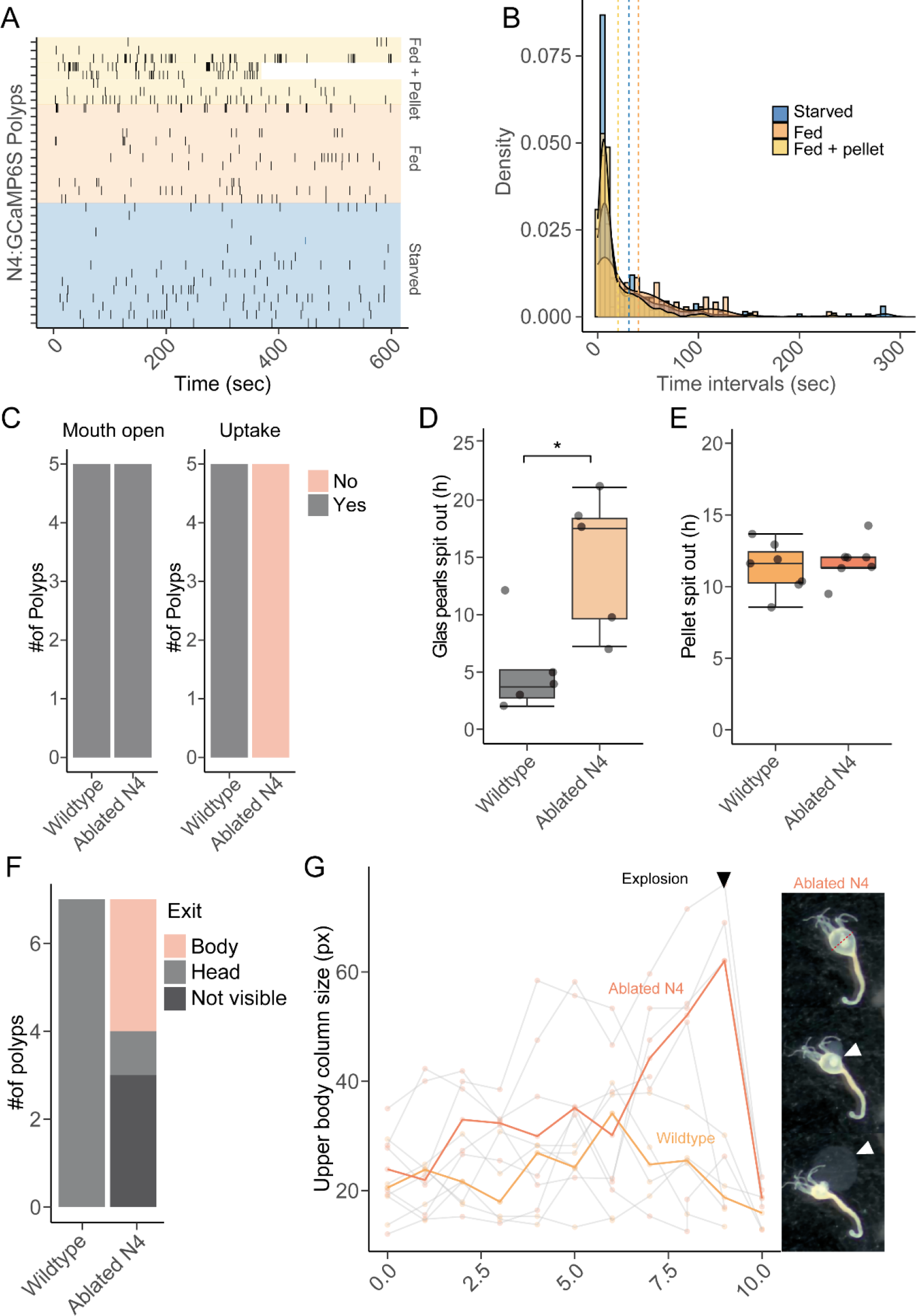
Endodermal N4 population regulates engulfment as well as spit out of digestive pellet. **A**. Spike train of endodermal population N4 4 hours post feeding with (yellowish) and without pellet (orange) and starved (blue). **B**. Inter spike interval distribution, fed with pellet has shorter intervals between consecutive spikes. **C**. Animals with ablated N4 neurons can open their mouth but cannot engulf glass beads (n = 5). **D**. Animals with ablated N4 neurons have difficulty to expel glass beads (n = 5, T-test). **E**. Time till spit out of artemia did not differ in N4 ablated and wildtype animals (n =6) **F**. The exit point of the digestive pellet varied between N4 ablated and wildtype. In N4 animals the pellet did not exclusively came out of the mouth (n = 6). **G**. After 6h post feeding, animals with ablated N4 population have become inflated underneath the head till they exploded which led to pellet exit through the through the ruptured tissue (n = 6, picture series).

To summarize our findings, we demonstrate here that *Hydra* has a long-lasting internal state which affects behavior and is controlled by two different neuronal networks. On one hand, the endodermal N4 network controls the response to the presence (or absence) of the digestive pellet and the ultimate spitting out of the pellet. On the other hand, the ectodermal N3 networks spiking frequency increases in a satiety dependent manner reducing the occurrence of other behaviors such as contractions and probably somersaults. Furthermore, N3 inhibits mouth opening when fed and ablation of N3 cancels the effect. These findings extend the recently described neuronal circuit of the eating behavior ^16^ and show that in *Hydra* functions of the digestive system and the control of behavior are coordinated by two very distinct neuronal populations.

## Discussion

Most animals adjust their behavior depending on whether they have ingested food or not. Often these behavioral changes are modulated by the brain which changes sensory processing based on the metabolic internal state and the enteric nervous system controlling food passage^1,19^. Astonishingly, internal states can already be observed in animals that have no centralized nervous system. The underlying neuronal control is unknown^5–8^. Here we show in *Hydra* that neuronal population N3 with a polyp-wide spanning network has a metabolic state dependent change in its activity and controls behavior as well as repeated food intake. In addition, the endodermal neuronal population N4 controls behavior associated with the feeding and digestion process, including the expulsion of the digestive pellet. These findings highlight a sub-functionalization within *Hydras* nervous system into an ectodermal network controlling motor and physiological functions and an endodermal network controlling feeding associated behaviors.

### Starved animals exhibit increased explorative behaviors

Behaviors associated with locomotion and mouth opening are reduced or even inhibited for more than 24 hours post feeding while contractions return much faster to a normal frequency (**Fig. 2A-F**). The exposure to glutathione in fed animals (food stimulus) did not result in a feeding response for more than 10 hours post feeding. Replacing the feeding with a pre-exposure to glutathione failed to reproduce the inhibiting effect for the same duration (**Fig. 2A**). This showed that the mere performance of the eating behavior could not prevent another feeding behavior 4h after the initial stimulus suggesting that the presence of real food is required to inhibit another response. A similar dependency was observed in locomotion associated behaviors. Somersaults were completely absent in fed animals for over 28h post feeding as well as fed animals did not detach from their substrate (**Fig. 2D, F**). Thoughts on the ecological relevance of the floating behavior of Hydra were already presented more than 60 years ago ^20^. It is to add, that fed animals were not motionless or devoid of activity, in fact they displayed higher levels of body movement compared to starved animals (**Fig. 2E**). However, fed animals are rather stationary whereas starved animals seem to do more locomotion, contractions and floating behavior.

### Neuronal control of digestion and internal state

The endodermal neuronal population N4 plays not only a crucial role in initiating the eating behavior of *Hydra* as previously shown^16^, but is also necessary for the downstream behavioral pattern. The N4 network displays an increased spiking frequency when a digestive pellet is present in the gastrovascular cavity, and this frequency is significantly reduced in the absence of the pellet (**Fig. 3D**). Further, cell ablation experiments demonstrate that N4 regulates the engulfment of the prey and the spit out of a digested pellet(**Fig. 5**). Both functions are supported by the presence of sensory cells in head of *Hydra*, extending into the gastrovascular cavity. These sensory cells are likely responsive to both chemical and mechanical stimuli (**Fig. 3F**). So far, N4 has not been associated with any other behavior and therefore we propose that N4 is mainly responsible for feeding-related behaviors and internal fluid transport.

The ectodermal neuronal population N3 is mainly involved in locomotion behavior and integrating physiological information such as the feeding state. The N3 network has previously been described to be involved in motor control of *Hydra* such as somersaulting and detachment^18^. Further, it counteracts the (contraction burst) N1 network^14,17,18,21^, is associated with elongation behavior, responds to blue light, osmolarity, and has non-behavior associated activity^18,21,22^. Here, we show that the activity of the N3 network increases and decreases in a feeding dependent manner and inhibits mouth opening while suppressing other behaviors such as body movement, phototaxis and somersaults (**Fig. 3G, Fig. 4**). The N3 neuronal network is unique due to its connectivity with the N1 network, as well as all other ectodermal neuronal populations and nematocytes, which allows for global modulation (**Fig. 3I**)^16,18^. Intriguingly, N3 neurons appear to be multi-functional as they are involved in various behavioral patterns each characterized by a unique spiking frequency (**Fig. 4D**)^17,18^. In the case of fed animals, the spiking pattern is strikingly regular and misses pattern such as bursts as seen in somersault behaviors^18^. This clearly shows that a single ectodermal neural population, population N3, possesses the ability to control numerous behaviors and to integrate comprehensive information about the organism’s condition. Our findings suggest that the rise in activity triggered by feeding is the result of modifications in the intrinsic neural activity, leading to substantial effects on adjacent circuits and the consequent regulation of behaviors. These observations indicate the existence of a hierarchical structure within the neuronal signaling system^14^.

Nevertheless, one question that has yet to be resolved is whether the alteration in activity of the N3 population can be attributed to endogenous regulation or exogenous influences. The activity of N3 has been associated to non-behavior associated activity (**Fig. 4D**)^17,23,24^. Here, we observed a significant higher spiking frequency in non-behavior associated activity in fed animals suggesting an intrinsic activation. In addition, the activity of N3 mainly emerges from the foot region and the activity appears to counteract the activity of N1^17,18^. Since there is no other ectodermal neuronal population that might trigger N3 and serve as the extrinsic driver, it is worth considering whether epithelial cells in the immediate proximity might not play an important role here. Perhaps subsequent work will provide clarity here.

In conclusion this work highlights the sub-functionalization of *Hydra’s* previously called “diffuse” nervous system into a subpopulation similar to the enteric nervous system (feeding associated behavior) and into a population that is performing central nervous system (physiology/motor) like functions. We further demonstrate that the ectodermal neuronal population changes its activity depending on the internal state and affects other behaviors. Taken together, *Hydra’s* nervous system serves as a remarkable illustration of how a simple nervous system can manifest intricate sub-functionalization; and perhaps for this reason it can be an important key to understanding the evolution of the Bilaterian nervous system.

## Acknowledgements

This work was supported in part by grants from the Deutsche Forschungsgemeinschaft (DFG), the CRC 1182 “Origin and Function of Metaorganisms” (to TCGB.) and the CRC 1461 “Neurotronics: Bio-Inspired Information Pathways” (Project-ID 434434223 – SFB 1461) (to TCGB). T.C.G.B. appreciates support from the Canadian Institute for Advanced Research. We thank the whole Bosch Lab and in particular Lisa-Marie Hofacker for support during the preparation of the manuscript. We also thank Urska Repnik and Marc Bramkamp from the Central Microscopy Facility at the Biology Department of the University of Kiel for excellent technical support.

## Authors contribution

C.G. and C.N conceptualized the project and conducted the experiments. C.G. generated the transgenic constructs and designed experiments for behavioral analysis. C.G. and C.N. performed cell ablation experiments. C.G., C.N., and E.S. performed calcium imaging experiments. C.G. and C.N. did data analysis. C.G., C.N. and T.C.G.B wrote manuscript.

## Data and code availability

- Source data reported in this paper will be shared by the lead contact upon request.
- Codes used for the analysis and statistical analysis will be shared by the lead contact upon request.
- Any additional information required to reanalyze the data in this paper will be shared by the lead contact upon request.

## Material and Methods

The plasmids and transgenic *Hydra vulgaris* AEP generated in this study are available upon request.

## Code availability

All codes used in this study are available upon request.

## Data availability

All data presented in this study are available upon request.

## Hydra maintenance

*Hydra* polyps (*Hydra vugaris* AEP) were subjected to cultivation procedures in standard *Hydra* culture medium (i.e., CaCl2 0.042g/L; MgSO4 x 7H20 0.081g/L; NaHCO3 0.042g/L; and K2CO3 0.011g/L in dH2O), in accordance with established protocols^25^. The animals were cultured in 250mL glass beakers maintained at a temperature of 18°C, with a 12/12h light cycle. Feeding of the animals strictly occurred three times per week, utilizing *Artemia nauplii* for at least two weeks prior to any experimentation.

### Behavioral analysis

#### Feeding behavior

To investigate the eating behavior and its dependency on the feeding state we followed a recently established protocol^16^. The animals were given a period of 10 minutes to adapt themselves to the recording chamber prior to the start of recording, followed by an additional interval of 5-10 minutes prior to the addition of reduced glutathione (Roth, cat#6382.1) through a tube system. A final concentration of 10µM glutathione was employed for all assays. The acquired recordings were blinded to their treatment and categorized with a random three-digit number before analysis. The conduct was manually annotated, with the following various behaviors being tallied (**Fig. 1E**): the duration of mouth opening and the response time. The duration of the mouth opening is the time the mouth stays open. The response time is the time an animal needs to show the first response to glutathione, here mouth opening.

#### Contractions, somersaults, body movement, and floating

Animals were placed into a custom-made chamber with a height of 5mm and a volume of 1.5-2mL which allowed the animals to move freely. Five animals were recorded at once with a frame rate of 0.5 frames per second. Further the recordings were blinded, and behaviors were manually annotated. For contraction behavior, all contractions were counted which reduced the animal length by a minimum of 33%. Somersaults were defined as the following sequence of behaviors: detachment of foot, attachment of head, contraction, bending and re-attachment of the foot in accordance with previously published work^18^. For the body movement, the body center of the animals was tracked by using Track Mate in ImageJ^26,27^ and the average speed (µm/sec) was calculated. For the floating behavior the lid of the chamber was removed which then allowed animals to flow on the surface. Here, single animals were tracked, and behavior was annotated as attached or detached/floating (**Fig. 1E**).

### GCaMP6S recording

In order to examine the neuronal activity in dependency of the feeding status, the animals were placed in commercially available channel slides that were 0.2 or 0.4mm in height and 5mm in width (Suppl. Fig. 6D.; Ibidi, cat#80166, cat#80176). Once an animal was placed in the channel slid, we led the animal adapt for 10 min to the new environment plus another 2min to the laser prior recording. Imaging was performed using the Axio Vert. A1 (Zeiss) with the Colibri 7 as a light source (Zeiss) equipped with the fluorescence filter 38 HE (Zeiss), 5x and 10x Plan Apo objective, and the Axiocam 705 mono (Zeiss). Acquired videos were further processed with Zen Blue 3.4 (Zeiss) to 700x600px, 8-bit and analyzed via either ImageJ or ICY^28^. The neurons were then tracked with a previously published pipeline^29^. The acquire tracks were then merged to one by taking the mean fluorescence change to analyze the population activity pattern.

### Neuronal activity analysis

The mean activity of each population or neuronal type was taken for further analysis with the standard deviation to the mean, as it summarized all major events. All visualization and normalization were performed using customized scripts in R^30^. The spiking frequency of N3 and N4 was deconvoluted and analyzed using either CASCADE^31^ and/or MATLAB’s (Mathworks) “findpeaks” function with manually adjusted parameters. At least 3 animals were used in all experiments.

### Ablation experiments

To test the role of population N3 and N4 in modulating the behavior post feeding, we used animals carrying the NTR-MTZ system under the control of a N3- or N4-specific promotor^16^. we fed the animals extensively and transferred them into 10mM metronidazole (MTZ) after 30min post feeding. For analyzing the behavioral changes, animals were placed in the costume made chambers as described under behavioral analysis. For testing the ability to engulf glass beads, we incubated N4::NTR-GFP animals overnight in 10mM MTZ together with a control (without a construct) and performed the mock feeding assay.

### Testing engulfment and spit-out (glass beads)

In order to test if animals lacking a certain neuronal population (N4) can still ingest prey and spit out the digestive pellet we force fed *Hydra* with glass beads with a diameter of 200-300µm (Sigma, #G9143). Animals were transferred into 10µM glutathione solution and then with a syringe or forceps stuffed with glass beads (>4<7 beads per animal). The ability to engulf a glass bead was scored. For testing the impact on the spit out time, animals were transferred into 10mM MTZ and recorded for 24h in the setup described under “Contraction, somersaults, body movement, and floating”. The time point where all glass beads were spit out was noted.

### Phototaxis assay

To investigate whether animals that have been adequately nourished (4-10 hours after feeding) exhibit alterations in their phototaxis behavior, and if these alterations are dependent on the N3 neuronal population, we conducted experiments using specially designed arenas. For this purpose, we utilized cell culture dishes with dimensions of 35×10mm (Eppendorf, cat#0030700112) and covered the walls with black tape, leaving a narrow opening measuring 0.5mm to allow light to enter the chamber. To provide the necessary light source, we employed a cold light source (Schott KL 1600 LED) with a high-power LED that emitted a maximum of 680 lumens and had a color temperature of approximately 5600 Kelvin. Prior to the commencement of the recordings, the animals were fed a substantial amount of *Artemia* until they were fully satiated and were then left at a temperature of 18°C for 30 minutes to initiate the digestion process. Following this 30-minute period, the animals were transferred into a solution of 10mM MTZ and were incubated for 4 hours in darkness at a temperature of 18°C. Subsequently, the animals were placed in the center of the arenas and were recorded for a duration of 24 hours at a temperature of 18°C. The recordings of the animals were captured at a frame rate of 0.5 frames per second, and the resulting movies were further processed using ImageJ^26^. The tracking of the animals was performed using TrackMate, and the log-threshold approach was utilized for this purpose^27^. The tracks obtained for each animal were subsequently subjected to additional processing and were visualized using R^30^. For comparison, the control group in all experiments consisted of wildtype polyps (*Hydra vulgaris* AEP) subjected to the same experimental conditions as the transgenic animals.

### Transgenic lines

The transgenic lines N4 (GCaMP6S and NTR-GFP) and N3 (GCaMP6S and NTR-GFP) have been generated in another study^16^. The pan-neuronal line was derived by isolating the promotor of a neuronal specific alpha tubulin (t14440aep/G019559, 1200bp) and this line has been already described^15^. Here, we derived this line to a F1 generation.

### Histology

Immunohistochemical detection was performed on whole-mount *Hydra* preparations using as previously described^16^. The polyps were relaxed (Urethane), fixed (Zamboni), permeabilized (0.5% Triton X-100 in PBS), incubated with primary antibodies (1:1000) and secondary antibodies (1:1000). Confocal laser-scanning microscopy was then used for analysis (LSM900, Zeiss). As primary antibody was used: chicken-anti-GFP (Biozol, cat# GFP-1010) and as secondary antibody: goat anti-chicken Alexa Fluor 488 (Invitrogen, cat# A11039, 1:1000 dilution).

### Data analysis

All statistical analyses were conducted using R and R-studio as the integrated development environment (IDE)^30,32^. In all instances, data were evaluated for equal variance using Levene’s test and normal distribution using the Shapiro test. Depending on the results of these tests, either a parametric (t-test, ANOVA, Turkey test) or non-parametric test (Kruskal-Wallis, pairwise or Wilcox test, Dunn test) was employed. Multiple testing correction was performed using Bonferroni. The number of replicates (n) for each dataset was indicated in the figure captions, along with the statistical method utilized for each comparison and the corresponding p-value. For linear correlation analysis, correlation coefficients and p-values were calculated using the Spearman method with the assistance of the R package “ggpubr”. The cutoff for a significant difference was established as an α < 0.05. Throughout the text, values are presented as median ± standard deviation, unless otherwise specified.

